# KymoTip: High-throughput Characterization of Tip-growth Dynamics in Plant Cells

**DOI:** 10.1101/2025.06.27.661917

**Authors:** Zichen Kang, Yusuke Kimata, Tomonobu Nonoyama, Toru Ikeuchi, Kazuyuki Kuchitsu, Satoru Tsugawa, Minako Ueda

## Abstract

Live imaging data analysis often requires an objective, local, and accurate way of quantification of cell dynamics. In the research field of polarized tip-growth, the cell fluctuations and/or fluctuations in tip position and growth direction hampers automated analyses of huge amounts of imaging sequences. The fluctuated nature in data makes it unclear how cell shape and growth are linked to intracellular events that could be the actual driving force of cell growth. To overcome these difficulties, we developed a powerful and user-friendly tool called KymoTip with an available format. In this software, novel functions such as coordinate normalization, tip-bottom detection, and signal kymograph were implemented. We confirmed that not only plasma membrane-labeled fluorescent images, but also images such as bright-field and cortical microtubule markers —so long as the cell contours can be identified— are amenable to KymoTip. Furthermore, by combining markers for cell contours with those that visualize intracellular structures, it becomes possible to quantitatively analyze various intracellular events, such as nuclear migration and calcium wave, in conjunction with cellular growth dynamics. Since KymoTip can be handled by non-specialist, it is expected to promote understanding of what happens at the sub- and cellular level with high throughput outcomes.

**Significance statement:** Faced with fluctuations in cell coordinates and cell tip positions, position correction of live imaging data and accurate detection of tip position are key challenges in plant developmental biology. We solved them with a powerful and user-friendly tool, KymoTip, that can realize cell position correction, cell tip detection with cell centerline, and quantification of intracellular events.

## Introduction

Tip growth dynamics is an important research topic from the viewpoint of developmental biology. For example, an anisotropic elongation of plant zygote in *Arabidopsis thaliana* (Arabidopsis) was recently found to be the type of tip-growth, indicating that the tip growth may be essentially important for understanding the first recognition of apical-basal axis for zygote in development (Kang et al., 2023). This finding facilitates obtaining different types of new live cell imaging so that huge amounts of data with high quality images and a well stored database including the wild type, mutants, and those with pharmaceutical treatment are ready to be analyzed (Hiromoto et al., 2023; Matsumoto et al., 2021; Kimata et al., 2023). These analyses revealed various polar dynamics, such as mitochondrial migration toward the growing cell (Kimata et al., 2020). However, due to the lack of a simple method capable of simultaneously capturing the continuously deforming cell outline and the dynamics of intracellular structures and events, quantitative analysis of intracellular dynamics in relation to cell growth still relies on static images and/or manual observation.

The main difficulty that hampers the data analysis was the cell fluctuation due to the experimental observation with the seeds in the liquid medium so that it needs a certain coordinate normalization method at the analyzing stage to fix the zygote cell contours. This positional fluctuation made it disabled to observe intracellular dynamics in a spatially rearranged manner. For other examples, the root hairs in Arabidopsis and the rhizoid in the liverwort *Marchantia polymorpha* (Marchantia) exhibited tip growth to anchor the plant body (Shaw et al., 2000, Otani 2018). The difficulty in this case was the fluctuation of the cell tip positions which often required researchers to stay focused and detect the tip manually as a first attempt so that an objective detection of the cell tip may break down the barriers. In these examples, the quantification and characterization of cytoplasmic signals relative to the growing tip of cells have also been required. However, automated data analysis of tip positional detection is challenging because the cell tip positions are often fluctuating.

In recent years, quantitative approaches that promote the analysis of the inner dynamics of plants and their interactions have profoundly influenced the field of plant biology (Autran et al., 2021; Hamant, 2024). In recent years, with the advancement of live-cell imaging techniques, the demand for quantitative image analysis that is capable of automated batch processing has been steadily increasing. For instance, PlantVis automatically analyzed the spatio-temporal behavior of root growth in Arabidopsis (Roberts et al., 2010; Wuyts et al., 2011). Similarly, RootflowRT quantified the growth rate of different types of root species (Weele et al., 2003), and KineRoot calculated growth rate without using characteristic markers for growth analysis (Basu et al., 2007). These software programs achieved objective detection of growth rate. The growing region can be detected by GrowthTracer, where the displacement of any positions of root can be quantified using a certain filter (Iwamoto et al., 2013). For extraction of the growth rate along the cell axis, KymoRod uses Voronoi tessellation to detect the cell centerline objectively (Bastien et al., 2016). Admitting that all these existing tools can quantify the growth rate or growth region of the plant organs, none of these techniques identify the cell tip due to the existence of cell fluctuations in position and in growth. Therefore, it has been required to develop an objective and accurate way of cell tip detection in plant cell biology.

In this study, we propose a novel user-friendly tool KymoTip with objective and accurate quantifications based on coordinate normalization. Firstly, we demonstrated a flow map of data analysis and concrete output of our analysis. Next, we showed the sufficiency of our method for quantitative analysis of cell growth dynamics using three examples: Arabidopsis zygotes, root hairs, and *Marchantia* rhizoids. Finally, we show the simultaneous analyses of cell growth dynamics and various intracellular events, and discuss how the objectivity and accuracy of tip growth are important for plant cell biology.

## Results and discussion

### 1. Algorithm and series of output by KymoTip

The algorithm to evaluate the tip-growth dynamics consists of three main steps. (1) Binarization/Segmentation to get the cell contours and their centerlines over time, (2) Coordinate normalization if necessary, and (3) Tip-bottom detection by extending the Voronoi tessellation.

At first, we obtained the cell segmentation and cell contours to get cell centerline from the original raw data, i.e., time-lapse images of Arabidopsis zygotes expressing green and red fluorescent markers labeling the plasma membrane and nucleus, respectively (Fig. 1A-C). We then obtained a skeletonization profile using Voronoi tessellation of the point cloud on the cell contour (Fig. 1D, see also Bastien et al., 2016). We then removed the small branching by detecting the intersections of skeletons and obtained the centerline that does not reach the contour (Fig. 1E). In these processes, we applied a method of segmentation based on SAM2 which enables a more accurate segmentation rather than a canonical method like Otsu (Ravi et al., 2024, see experimental procedures, Fig. 1F). Second, we applied coordinate normalization detailed in the experimental procedures (Fig. 1G). Third, we detected the cell tip and bottom by extending the centerline (Fig. 1H).

**Fig. 1.**
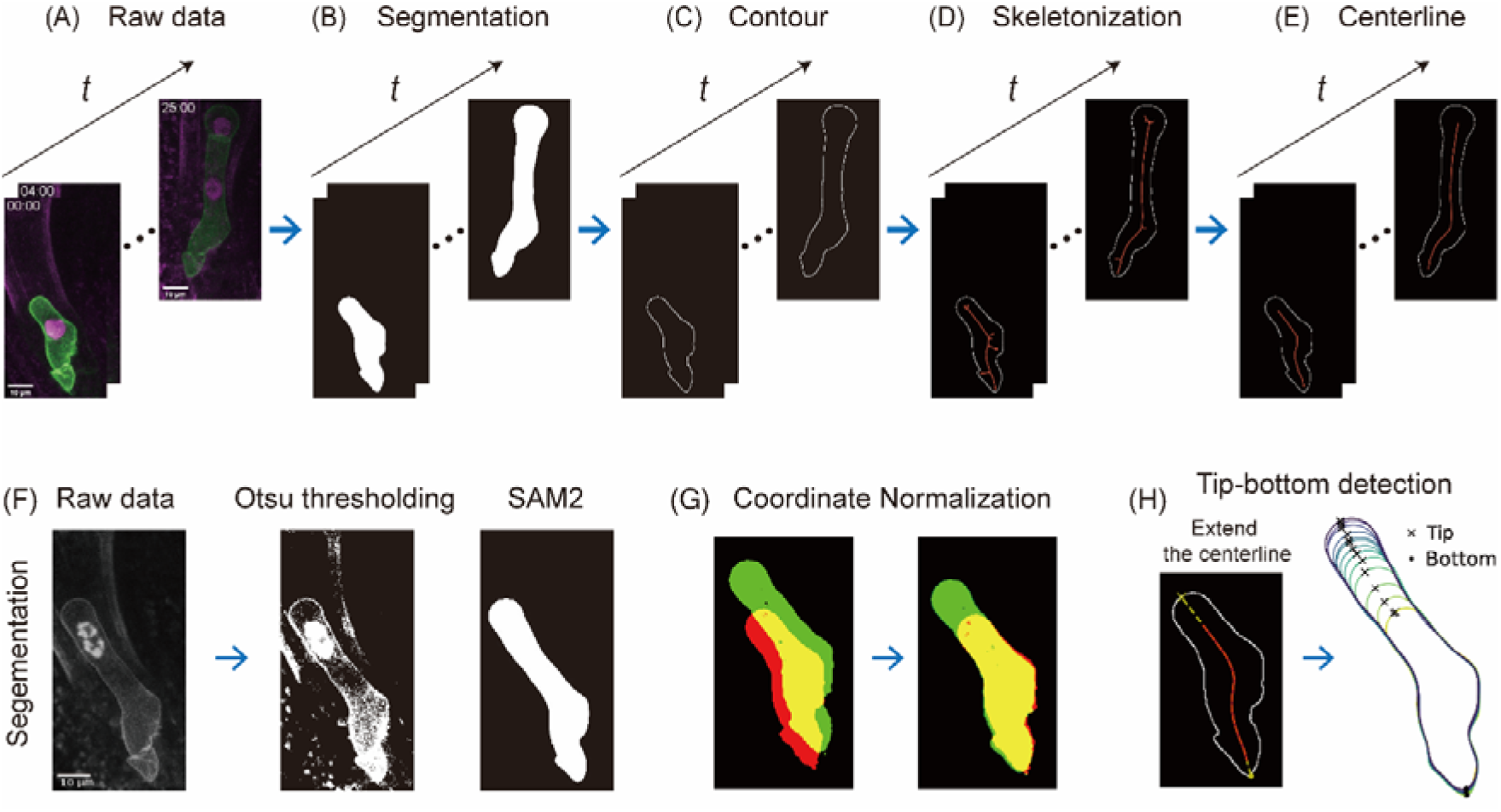
Workflow of KymoTip. (A) Raw data. Scale bars, 10μm (B) Binarization/Segmentation. (C) Cell contours. (D) Skeletonization using Voronoi tessellation. (E) Cell centerline. (F) Comparison between Otsu thresholding and SAM2. (G) Coordinate normalization of two successive images. (H) Tip-bottom detection by extending the centerline to the cell edge.

### 2. Applicability of KymoTip to various tip-growing cells

We propose new methods of detecting the cell tip position, velocity and direction. We put three examples of tip detected results for zygote, root hair and rhizoid (Fig. 2A). Using tip positions, we could calculate the tip velocity dL/dt with the cell length L and the velocity direction Δθ as schematically illustrated in Fig. 2B. Using tip positions of zygote (Fig. 2C left), we obtained that the tip velocity was maximally 0.05 um/min and the velocity direction indicates that the tip first faces to the left gradually changed the direction to the right in this example (Fig. 2D). In the case of root hair of time-lapse bright-field images (Fig. 2C center), the cell centerline and tip detection were performed without the coordinate normalization. We note that when the cell bottom was fixed experimentally the coordinate normalization is not necessary. Then, we obtained that the tip velocity was maximally 0.2 µm/min roughly estimated by 4 times faster than zygote in which the velocity direction is not significantly changed (Fig. 2E). From raw data of *mCherry* fluorescence in the rhizoid (method), it shows bumpy cell shapes over time (Fig. 2C right). The maximal tip velocity was 0.6 um/min that was roughly three times faster than root hairs (Fig. 2F). More interestingly, when the tip velocity becomes higher, the velocity direction is also changed synchronously.

**Fig. 2.**
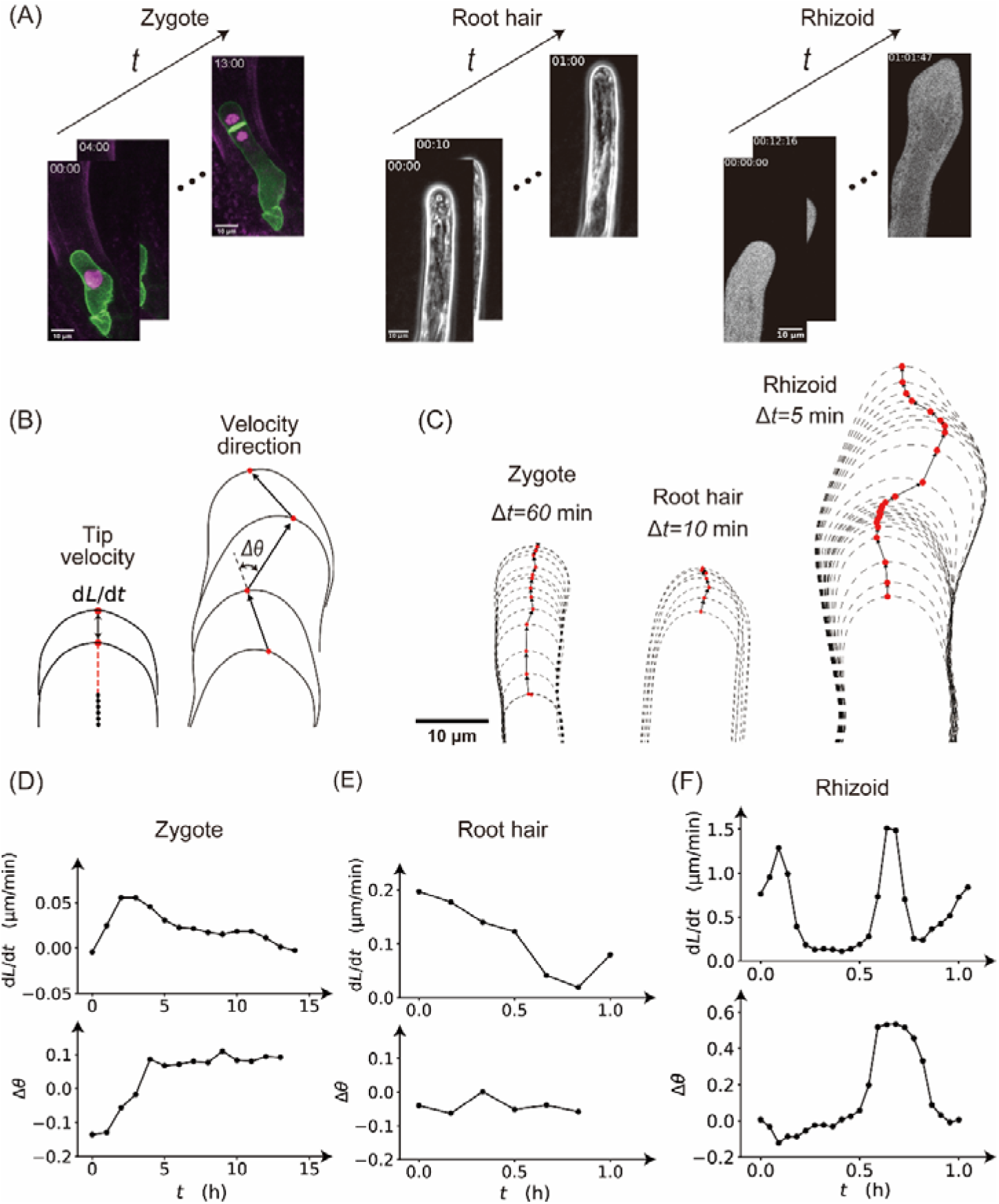
Versatility of the KymoTip of detections of the tip-bottom, tip velocity and velocity detection. (A) Raw data of zygote, root hair, and rhizoid. (B) Schematic illustrations of the tip velocity dL/dt and the velocity direction Δθ by tracking the cell tip over time. (C) Examples of tip detections of zygote, root hair, and rhizoid. Δt denotes the time interval of the image sequences. (D-F) Quantitative results of tip velocity and velocity direction of zygote (D), root hair (E), and rhizoid (F).

### 3. Quantification of cytoplasmic signals

We further investigated whether KymoTip enables simultaneous tracking of intracellular events in relation to changes in cell morphology. For this purpose, we examined the spatiotemporal dynamics of nuclear migration toward the apical cell tip and the constant accumulation of a microtubule band in the subapical region of the zygote (Kimata et al, 2016; Kang et al 2024).

First, the nuclear position was determined using dual-color imaging of the cell outline and nuclear markers (as in Fig. 1A). We applied KymoTip to extract the cell centerline from the cell contour (Fig. 1), and fitted the fluorescence profile of the nuclear signal using a truncated Gaussian distribution (Fig. 3A). Based on these data, a kymograph was generated (Fig. 3B), allowing simultaneous quantification of both cell growth and nuclear movement (Fig. 3C). These results demonstrate that various intracellular dynamics can be precisely quantified by converting them into a well-defined coordinate system along the cell centerline, enabled by co-visualization with a cell contour marker. Furthermore, when using live imaging data of microtubule markers, the cell shape was inherently visualized by the marker itself (Fig. 3D). Thus, even without an additional cell contour marker, KymoTip successfully enabled the tracking of microtubule position in the cell coordinate system (Fig. 3D–F). In this context, the constant microtubule band in the subapical region was reliably detected using truncated Gaussian fitting (Fig. 3E and 3F). These features highlight the significant advantages of KymoTip.

**Fig. 3.**
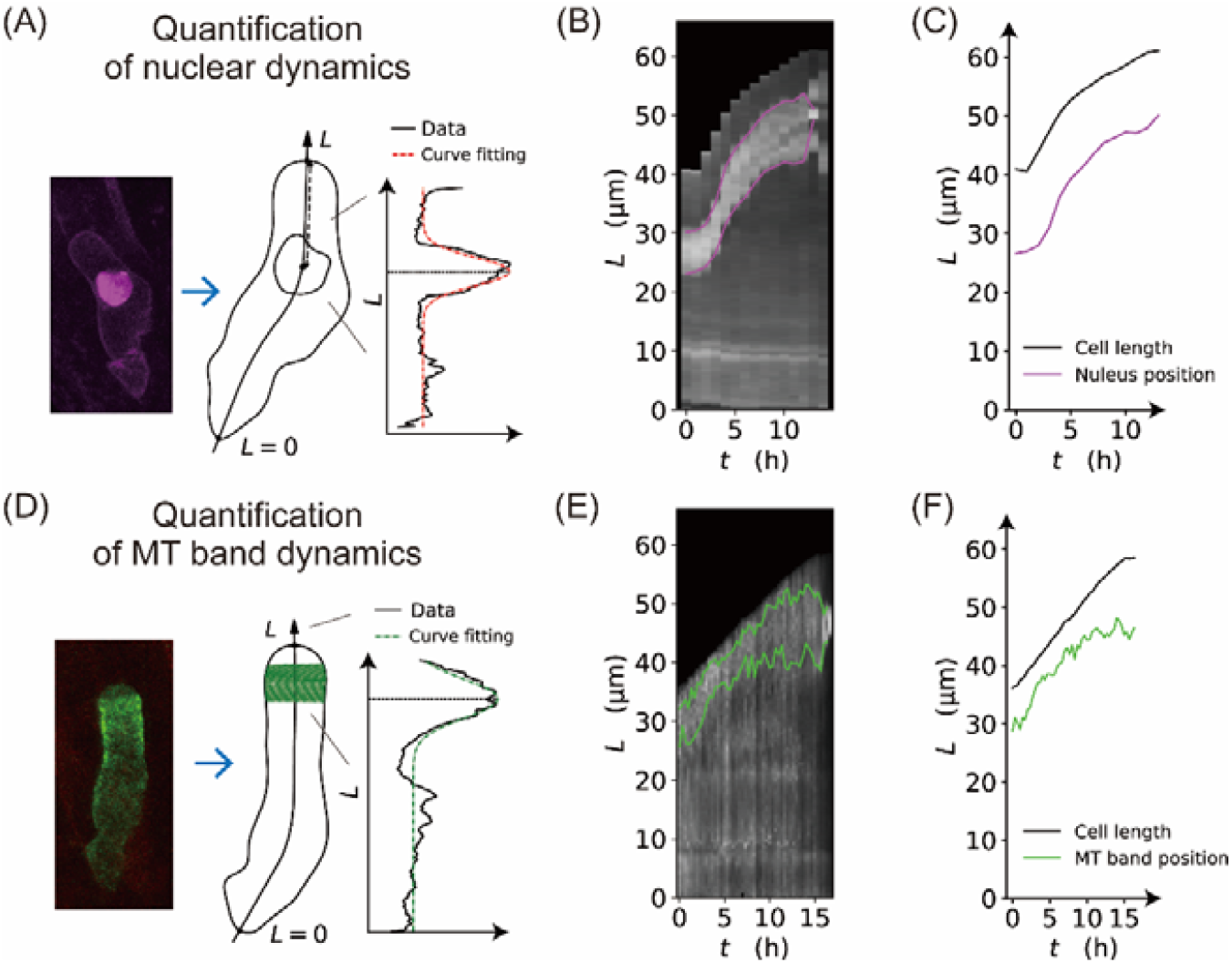
(A) Quantification of nuclear dynamics in zygote by fitting the curve of the nucleus fluorescence intensity (Kang et al., 2024). (B) Kymograph of nucleus fluorescence based on coordinate normalization in zygote. Magenta lines show upper and lower ends of the nucleus. (C) Quantitative comparison of cell length and nucleus position in zygote. (D) Quantification of microtubule band by Gaussian fitting of the microtubule fluorescence intensity in zygote (Kang et al., 2024BioRxiv) (E) Kymograph of microtubule band based on coordinate normalization in zygote. Green lines show upper and lower ends of the microtubule band. (F) Quantitative comparison of cell length and microtubule band position.

### 4. Summary and future perspectives

In summary, we developed a powerful tool KymoTip to assess tip growth dynamics, allowing improved segmentation with SAM2, coordinate normalization, tip detection by centerline extraction, quantification of growth direction, and an evaluation of the spatial distribution of cytoplasmic signals in plant cells. The first highlight was that an improved cell contour segmentation of SAM2 resulted in more accurate precision of coordinate normalization, enabling the fixation of cell fluctuations. The second highlight was that an idea of extension of the centerline leads to an accurate detection of the cell tip, enabling the quantification of growth direction and spatial cytoplasmic signals. The method is available on the internet which prompts understanding of various tip-growth dynamics for ubiquitous users.

For future perspectives, three novel properties will be available to analyze in tip growing dynamics: cell contours with coordinate normalization, tip positions, and its growth directions. With respect to the normalized cell contours, it allows ones to clearly see the cytoplasmic fluorescence being freed from visual fluctuations of cell position. In addition, a spatio-temporal curvature change of cell shape can be traceable in a well rearranged manner. With respect to tip positions, a dream of visualizing the cytoplasmic signals centering from the cell tip comes true. Furthermore, a relative distance of cytoskeleton signals such as microtubule band to the cell tip can be quantified, enabling one to think of its biological meaning during tip growth. With respect to tip growth directions, it may be able to see correlations between the calcium waves and growth direction, and it may also be able to understand the relationship between growth rate and growth direction. Importantly, the KymoTip method is not limited to a specific plant species or cell type. When combined with our recently developed technique that enables the visualization of the living zygotes across a wide range of plant species using only fluorescent staining (Hanaki et al., 2025), KymoTip offers a general framework for high-resolution and quantitative analysis of cellular dynamics beyond conventional model organisms. Thus, KymoTip definitely opens a new avenue for understanding multi-faceted aspects of tip growth and cellular behavior across plant lineages.

## EXPERIMENTAL PROCEDURES

### Image acquisition and pre-treatment

Live imaging data of the plasma membrane and nucleus in Arabidopsis zygotes were obtained using an *in vitro* ovule cultivation method combined with two-photon excitation microscopy, and were used in a previous study (Ueda et al., 2020; Kang et al., 2023). Time-lapse imaging data of zygotic microtubules and root hair morphology have also been previously reported (Kang et al., 2023; Kang et al., 2024).

For imaging of *Marchantia polymorpha* rhizoids, the wild type (Tak-1) plants expressing mCherry were grown as reported previously (Watanabe et al., 2024). Gemmae were floated in 10 mL of MES buffer (0.5 mM CaCl□, 50 mM sorbitol, 0.25% low melting point agar, pH 5.7) and incubated for more than 16 h under continuous white light. For imaging, several gemmae were transferred to a drop of 10 mM MES buffer placed at the center of a coverslip bordered on all sides with vinyl tape, then sealed with a second coverslip to form a shallow chamber. Time-lapse imaging of mCherry fluorescence (excited by 577 nm laser and detected 561–628 nm) was performed by a confocal laser scanning microscope LSM900 (Carl Zeiss).

### Algorithms of the detections of contour and cell centerline

We used a deep-learning based model SAM2 (Ravi et al., 2024) for segmentation of the images. In this process, we need to identify the cell as an object by pointing to a specific point inside the cell. Using the points of the resulting cell contour, we applied the Voronoi tessellation associated with xy-coordinates of the cell contour to detect a skeletonization of the cell. Then, the centerline was obtained by cutting the small branches. We used x points near the end of the branch-cut skeletonization and extrapolated them linearly. We can use Lowess for cell contours and/or the resulting centerline to smooth the curves.

### Coordinate normalization

We rearranged the cell contours using an image-based coordinate normalization (Kang et al., 2023). We define *f*_ref_ (*x, y*) as the fluorescence intensity of the reference image at pixel coordinates (*x, y*), and *f*(*x*(*θ*), *y*(*θ*)) as the fluorescence intensity of the rotated image at the corresponding coordinates determined by the rotation angle *θ*. The variables *u, v* are the translational shift in *x*-and *y*-directions, respectively. We then calculated a normalized correlation function written as,

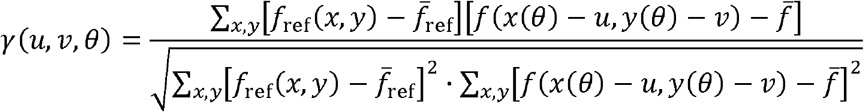

Here, 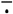 is the spatial average of. We note that the values of *f* (*x*(*θ*) − *u, y*(*θ*) − *v*) are set to 0 in regions that fall outside the image boundaries after rotation and/or translation. We optimized the parameters *u, v* and *θ* to maximize the correlation function and thereby align the image sequences with the reference frame.

## Code availability

The code for KymoTip is available on GitHub: https://github.com/blues0910/KymoTip. And the code for SAM2 segmentation is available at https://github.com/YusukeKimata-Moo/SAM2-segmentation/.

## Acknowledgments

This work was supported by the Japan Society for the Promotion of Science [a Grant-in-Aid for Early-Career Scientists (JP25K18499 to Z.K., JP23K14204 to Y.K.), a Grant-in-Aid for Scientific Research (B) (JP23H02494 to M.U., JP23H01143 to S.T.), Grant-in-Aid for Transformative Research Areas (A) (JP25H01809 to Y.K.)), International Leading Research (JP22K21352 to M.U.)], the Japan Science and Technology Agency [CREST (JPMJCR2121 to M.U. and S.T.) and Young Researcher Challenge (YORC) to Z.K. and T.N.)], the Suntory Rising Stars Encouragement Program in Life Sciences (SunRiSE; to M.U.), and the Toray Science Foundation (20-6102 to M.U.).

